# Dynamin-2 Mutations Linked to Neonatal-onset Centronuclear Myopathy impair exocytosis and endocytosis in adrenal chromaffin cells

**DOI:** 10.1101/2024.03.07.583689

**Authors:** Lucas Bayonés, María José Guerra-Fernández, Cindel Figueroa-Cares, Luciana I. Gallo, Samuel Alfonso-Bueno, Ximena Báez-Matus, Arlek González-Jamett, Ana M. Cárdenas, Fernando D. Marengo

**Affiliations:** Instituto de Fisiología, Biología Molecular y Neurociencias. CONICET. Departamento de Fisiología y Biología Molecular y Celular. Facultad de Ciencias Exactas y Naturales. Universidad de Buenos Aires, Buenos Aires, 1428, Argentina; Centro Interdisciplinario de Neurociencia de Valparaíso, Facultad de Ciencias, Universidad de Valparaíso, Gran Bretaña 1111, Valparaíso, 2360102, Chile; Escuela de Química y Farmacia, Facultad de Farmacia, Universidad de Valparaíso, Valparaíso 2360102, Chile

## Abstract

Dynamins are large GTPases whose primary function is to catalyze membrane scission during endocytosis, but also modulate other cellular processes, such as actin polymerization and vesicle trafficking. Recently, we reported that centronuclear myopathy associated dynamin-2 mutations, p.A618T and p.S619L, impair Ca^2+^-induced exocytosis of GLUT4 containing vesicles in immortalized human myoblasts. As exocytosis and endocytosis occur within rapid timescales, here we applied high-temporal resolution techniques, such as patch-clamp capacitance measurements and carbon-fiber amperometry to assess the effects of these mutations on these two cellular processes using bovine chromaffin cells as a study model. We found that the expression of any of these dynamin-2 mutants inhibits a dynamin and F-actin dependent form of fast endocytosis triggered by single action potential stimulus, as well as inhibits a slow compensatory endocytosis induced by 500 ms square depolarization. Both dynamin-2 mutants further reduced the exocytosis induced by 500 ms depolarizations, and the frequency of release events and the recruitment of NPY-labelled vesicles to the cell cortex after stimulation of nicotinic acetylcholine receptors with DMPP. They also provoked a significant decrease in the Ca^2+^-induced formation of new actin filaments in permeabilized chromaffin cells. In summary, our results indicate that the CNM-linked p.A618T and p.S619L mutations in dynamin-2 affect exocytosis and endocytosis, being the disruption of F-actin a possible explanation for these results. These impaired cellular processes might underlie the pathogenic mechanisms associated with these mutations.

## INTRODUCTION

Dynamins are a family of large GTPases that form high-order homo-oligomers and play pivotal roles in various cellular processes. These proteins consist of distinct domains, including a GTPase domain, a stalk region involved in self-assembly, a bundle signaling element (BSE) module that connects the G-domain with the stalk and transmits conformational changes upon GTP hydrolysis, a pleckstrin homology domain (PH) for membrane binding, and a proline-rich domain (PRD) (Arriagada-Diaz et al., 2020). Their primary function involves catalyzing membrane scission during clathrin-mediated endocytosis, caveolae internalization, and membrane budding (Arriagada-Diaz et al., 2020). Furthermore, dynamins are key regulators of cytoskeletal actin remodeling, contributing to diverse mechanical actions during cellular processes, such as bulk endocytosis, phagosome formation, neuronal growth cone dynamics, membrane protrusion, and fusion pore regulation (Yamada et al., 2016a; Chuang et al., 2019; Zhang et al., 2020; González-Jamett et al., 2013a; Shin et al., 2018; Linadu and Álvarez de Toledo et al., 2003).

Among the three dynamin isoforms expressed in mammals, dynamin-2 stands out as the ubiquitous variant (González-Jamett et al., 2014). Mutations in the gene encoding dynamin- 2 have been associated with two distinct neurological disorders, centronuclear myopathy (CNM) and Charcot-Marie-Tooth neuropathy (González-Jamett et al., 2013b; Ge et al., 2016). Notably, Charcot-Marie-Tooth-associated mutations predominantly affect the lipid-binding pocket within the PH domain’s N-terminal region, impairing dynamin’s affinity for lipids and disrupting clathrin-mediated endocytosis (González-Jamett et al., 2014; Koutsopoulos et al., 2011; Sidiropoulos et al., 2012). Conversely, CNM-associated mutations cluster in the middle domain, a region critical for oligomerization, or at the interface between the stalk region and the PH domain that coordinates lipid binding and GTPase activity (Ramachandran et al., 2007; Kenniston and Lemmon, 2010). These mutations often enhance dynamin-2 self-assembly, resulting in stable oligomers and heightened basal GTPase activity (Wang et al., 2010; Chin et al., 2015). Particularly noteworthy, the p.A618T and p.S619L mutations, located in the C-terminal α-helix motif of the PH domain, enhance lipid-stimulated GTPase activity and basal GTPase activity, respectively (Kenniston and Lemmon, 2010). Strikingly, these two mutations have been linked to severe neonatal forms of CNM (Böhm et al., 2012).

Dynamin-2’s canonical function lies in endocytosis, a process occurring within rapid timescales, necessitating high-temporal resolution techniques for accurate assessment (Chanaday and Kavalali, 2018; Montenegro et al, 2021). Therefore, to investigate the impact of p.A618T and p.S619L mutations on endocytosis, we employed capacitance measurements in patch-clamped adrenal chromaffin cells. Our findings revealed that both mutants significantly diminished fast dynamin-dependent endocytosis induced by short-term depolarization stimuli and the slow compensatory endocytosis induced by 500 ms square depolarization. Moreover, these mutations impaired the exocytosis triggered by 500 ms depolarizations, and the release events and recruitment of NPY-labelled vesicles to the cell cortex induced by stimulation of nicotinic acetylcholine receptors (nAChRs) with the agonist 1,1-dimethyl-4-phenyl piperazine iodide (DMPP). Notably, both mutations caused substantial disruptions of F-actin formation, as visualized through confocal microscopy of fluorescently-tagged G-actin. Consequently, both CNM-associated mutations adversely affected exocytosis and endocytosis, and these processes can be potentially attributed to dysregulated cytoskeletal actin filaments. These aberrant cellular processes might underlie the pathogenic mechanisms associated with these mutations.

## MATERIAL AND METHODS

### Plasmids

Wild type (WT) dynamin-2 fused to enhanced green fluorescent protein (EGFP) N1 rat was kindly provided by Dr. Stephane Gasman (CNRS, Strasbourg, France). The mCherry WT-dynamin-2 was constructed by GenScript Corporation (Nanjing, China) by cloning the rat dynamin-2 isoform aa (Genbank L25605) into an mCherry_pcDNA3.1(+) vector. The EGFP and mCherry dynamin-2 mutations A618T and S619L were constructed by site-directed mutagenesis using the QuikChange II XL Site-Directed Mutagenesis Kit (Agilent Technologies, Santa Clara, CA, USA). Neuropeptide Y fused to mCherry (NPY-mCherry) was kindly provided by Sebastian Barg (Gandasi et al, 2015, doi: 10.1371/journal.pone.0127801)

### Chromaffin cell culture and transfections

Bovine adrenal glands were acquired from local slaughterhouses (Frigorífico Cocarsa- Ecocarnes S.A., San Fernando, Provincia de Buenos Aires, Argentina, and Frigorífico Don Pedro, Quilpué, Chile). The isolation and culture of chromaffin cells was performed as previously published (Guerra et al., 2019). In brief, after adrenal medulla digestion with a solution containing 0.25% collagenase B, 0.01% trypsin inhibitor and 0.5% bovine serum albumin, chromaffin cells were isolated by Percoll gradient centrifugation and resuspended in a 1:1 mixture of Dulbecco’s Modified Eagle’s Medium/Nutrient Ham’s Mixture supplemented with 10% fetal bovine serum. The transfection of chromaffin cells with the plasmids described above was performed by electroporation in an Amaxa Nucleofector 4D (Lonza, Cologne, Germany) according to the manufacturer’s instructions. The cells were cultured at a density of 3 × 10^5^ cells/ml on 0.01% poly-l-lysine treated coverslips and kept at 37°C in a 5% CO2 atmosphere. Cells were used between 2 days after transfections.

### Whole Cell Patch-Clamp and Membrane Capacitance Measurements

These experiments were performed in an inverted microscope (Olympus IX71, Olympus, Japan), using a patch-clamp amplifier (EPC10 double patch clamp amplifier, Heka Elektronik, Lambrecht, Germany). The application of the stimulation protocols and the acquisition of data were controlled by the Patchmaster® software (Heka, Lambrecht, Germany, RRID:SCR_000034). The standard extracellular solution was composed of (mM) 120 NaCl, 20 Hepes, 4 MgCl_2_, 5 CaCl_2_, 5 mg/ml glucose and 1 µM tetrodotoxin (pH 7.3). The patch-clamp electrodes (3–5 MΩ) were coated with dental wax, and then filled with an internal solution containing (mM) 95 Cs D-glutamate, 23 HEPES, 30 CsCl, 8 NaCl, 1 MgCl_2_, 2 Mg-ATP, 0.3 Li-GTP, and 0.5 Cs-EGTA (pH 7.2). The holding potential of -80 mV was not corrected for junction potentials, and the recorded cells were discarded when the leak current measured at this holding potential was bigger than −10 pA, or when series resistance was bigger than 12 MΩ. The cell capacitance (Cm) was measured continuously during the experiments by application of the sine+dc method (Gillis 1995) implemented through the lock-in extension of Patchmaster, using a sinusoidal voltage (800 Hz, 60 mV peak to peak) added to the holding potential. The application of the sinusoidal voltage was suspended from 10 ms before to 10 ms after the end of the depolarizations applied to stimulate the cells.

The exocytosis was induced by a single 5 ms square depolarizations from the holding potential of -80 to -10 mV, or by an action potential-like stimulus (APls), composed by a 2.5 ms ascendant voltage ramp from -80 to +50 mV, followed by a 2.5 ms descendant ramp. In some cells, immediately after the application of a depolarization pulse, we noted the presence of a brief transient capacitance change, probably associated to sodium channels gating (Horrigan & Bookman 1994). This transient became negligible in less than 50 ms after the end of depolarization (Moya-Diaz *et al*. 2016; Chow et al. 1996; Gong et al. 2005). Therefore, to avoid any influence of this fast capacitance transient, the synchronous increase in Cm (ΔCm_exo_) associated with exocytosis was defined as the difference between the averaged Cm measured in a 50 ms window starting 60 ms after the end of the stimulus minus the average pre-stimulus Cm also measured in a 50 ms window (see the blue segments in Supplementary Figure 1). In a similar way, the following decreased in capacitance (ΔCm_endo_), associated with endocytosis, was calculated as the difference between the averaged Cm measured in a 50 ms window starting 60 ms after the end of the stimulus minus the averaged Cm measured 5 s after the stimulus, also in a 50 ms window (see the blue segments in Supplementary Figure 1). The Cm decay associated with endocytosis was fitted in each experiment to a single exponential function with the form 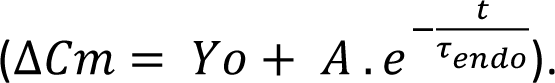. The data were filtered at 3 kHz. The experiments were carried out at room temperature (22–24◦C).

### Amperometric Recordings

Catecholamine release events were measured by amperometry as previously described (González-Jamett et al., 2017) using 5 μm diameter carbon fibers (Thornel P-55; Amoco Performance Product, Greenville, USA) which were held at a potential of +650 mV with HEKA EPC10 amplifier (HEKA Elektronik, Lambrecht/Pfalz, Germany). Cells were maintained in Krebs-HEPES solution (in mM: 140 NaCl, 5.9 KCl, 1.2 MgCl2, 2 CaCl2, 10 HEPES-NaOH, 10 glucose, pH 7.4) throughout the recordings, and exocytosis was induced with a 10 s of the nicotinic agonist dimethylphenylpiperazinium (DMPP) at 50 µM by using a custom-made picospritzer using a pressure of 2-3 psi. Amperometric signals were low pass filtered at 1 kHz and digitized at 5 kHz with the HEKA EPC10 amplifier and the PatchMaster software (HEKA Elektronik, Lambrecht/Pfalz, Germany). Each recording lasted 100 s and was analyzed using a custom-written macro for IGOR PRO (Wavemetrics Inc., Lake Oswego, Oregon, USA) gratefully obtained from Dr. R. Borges (Universidad de Tenerife, Spain). Analyses were restricted to spikes with amplitudes ≥ 10 pA, foot amplitudes ≥ 3 pA and foot durations ≥ 3 ms.

### Expression of Neuropeptide Y fused to mCherry as vesicle marker

Chromaffin cells were co-transfected with the construct NPY-mCherry together with Dyn-2 WT, or alternatively Dyn-2 p.A618T or Dyn-2 p.S619L. After 48 hours of transfection, cells were washed (x3) with PBS buffer, where one set of cells served as an unstimulated control, while the other was stimulated with the agonist DMPP (50 μM, 10 s). Then, the cells were fixed in a PBS solution containing 4% saccharose. After 24 h, cells were washed again with PBS and incubated for 10 min with PBS containing tris(hydroxymethyl)aminomethane (10 mM) to deactivate PFA innate fluorescence. Finally, after a final wash (PBS x5), cells were covered with montage liquid Mowiol® and storaged at 4°C.

Confocal images were obtained with a FV300 (OLYMPUS) confocal microscopy, using a Plan-Apochromat 60x/1.4 Oil DIC objective (OLYMPUS). To visualize the green-fluorescent marker EGFP, excitation was performed at 488 nm and the emission recorded at 500-520 nm; and for the mCherry red-fluorescent marker, excitation was applied at 543 nm and emission was collected at 600-620 nm. Images were acquired in X, Y and Z axes. In the Z axis, Z-stacks were used, with stacks being separated 0.35 μm from each other.

Images were analyzed in the equatorial plane by using a custom macro on Image J software (NIH, USA). The macro identified masks which accounted for the total and peripheral (1 μm width aligned with the border) areas, using as reference the area occupied by the fluorescence emitted for the EGPF-Dyn-2. Then, a second mask identified the area occupied by the fluorescence of the NPY-mCherry. Percentages of total and peripheral NPY were computed per cell as a ratio between the area associated to NPY-mCherry fluorescence and the celĺs total or peripheral area, respectively.

### Actin filament formation

Formation of actin filaments was carried out as previously described (Olivares et al., 2014). In brief, cultured cells were incubated in a buffer containing 139 mM k- glutamate, 20 mM PIPES, 5 mM EGTA, 2 mM ATP-Mg^2+^, 20 μM digitonin, 10 μM free Ca^2+^ and 0.3 μM Alexa Fluor 488-G-actin conjugate, at pH 6.6 with. Then, cells were fixed with 4% p-formaldehyde in phosphate buffered saline (pH 7.4), stained with 5 mg/ml 4′,6-diamidino-2-phenylindole and visualized using a Nikon C1 Plus laser-scanning confocal microscope (Nikon, Tokyo, Japan). All images were captured at the equatorial plane of the cells, using identical exposure settings between compared samples. Confocal images were analyzed and processed using a custom macro on ImageJ software (NIH, USA). The macro identified masks which accounted for: 1) the total celĺs area, using as reference the area associated to mCherry-Dyn-2 fluorescence, determined by Otsu’s thresholding algorithm; and 2) the area occupied by Alexa Fluor 488 fluorescence conjugate above a threshold determined by Renyís entropy based algorithm, a common standardized technique supported by ImageJ. Finally, the percentage of total F-actin formation was computed per cell as a ratio between Alexa Fluor 488 and mCherry-Dyn-2 associated areas.

### Statistics analysis

For every experimental condition, the data (one measurement per cell, and one cell per petry dish) were obtained from several cell cultures, which number is mentioned in every experimental condition. No blinding was performed in this study, and no randomization procedures were applied. No test for outliers was performed. Normality was assessed with Shapiro-Wilk and D’Agostino test. Normally distributed data were analyzed by Student’s “t” test (two-tailed) for comparisons between two groups of independent data samples, and by one way ANOVA (two-tailed test) for multiple independent data samples. After ANOVA, Bonferroni test (one-tailed) was used for comparisons between groups. Non-normally distributed data were analyzed using the Mann-Whitney test, or the Kruskal-Wallis test followed by Dunn’s post hoc test when pertinent. All statistical analyses were performed using Prism software version (GraphPad V5 Software Inc., La Jolla, CA, USA). To fit endocytosis records we used the nonlinear curve-fitting option in Origin Pro 8 (Microcal Software, RRID:SCR_002815). Data are expressed as means ± SEM for normal distributions. For non-normally distributed data, box plots were used showing a centered, inferior and superior lines, representing the median, first and third quartile, respectively. Top and bottom whiskers lines show the maximum and minimum values of each distribution, respectively.

### Materials

ATP-Mg (CAT No: A9187-100MG), Bovine serum albumin (CAT No. 05470-25G), CaCl_2_ (Cat No: 1.02382.1000), cytosine-1-beta-D-arabinofuranoside (CAT No: C6645-25MG), 4′,6- diamidino-2-phenylindole (DAPI) (CAT No: D9542), Digitonin (CAT No: D141-100MG), 1,1- Dimethylphenyl-4-phenilpiperazinium iodide (DMPP) (CAT No: D5891), EGTA (CAT No: E4378-10G), Glucose (Cat No: 1.08337.1000), GTP-Li (CAT No: G5884-25MG), KCl (CAT No: 1.04936.1000), MgCl2 (CAT No: 1.05833.0250), Mowiol® 4-88 (CAT No: 81381-50G), NaCl (CAT No: 1.0604.1000), P-formaldehyde in phosphate (CAT No: 158127-3KG), PIPES (CAT No: P-6757), and poly-L-lysine (CAT No: P8920-100ML) were obtained from Sigma-Aldrich (St. Louis MO USA); Dulbecco’s modified Eagle’s medium (CAT No: 10567-014), Fetal bovine serum (CAT No:16000-044), gentamycin (CAT No: 15750078), trypsin inhibitor (CAT No: R007100), and penicillin/streptomycin (CAT No: 15140122) from Gibco (Grand Island, NY USA); tetrodotoxin citrate (CAT No. T-550) from Alomone Labs (Jerusalem, Israel); Collagenase B (CAT No: 11088815001) from Roche; CsCl (CAT No: 102039) from Supelco - Merck (Darmstadt, Alemania); monoclonal anti-dynamin antibody (CAT No: 610246, RRID:AB_397641) from BD Biosciences (San Jose, CA, EEUU); Percoll (CAT No: 17-0891-02) from GE Healthcare (Uppsala, Sweden), and Alexa Fluor 488-G-actin (CAT No: A12373) from Thermo Fisher Scientific (Waltham, MA, USA)

### Ethics statement

All research undertaken had the approval of Bioethics (identification numbers BEA080 of Universidad de Valparaíso, Chile; and protocol 168 of the Facultad de Ciencias Exactas y Naturales, Universidad de Buenos Aires). This investigation comprises the use of bovine adrenal glands that were attained from local slaughterhouses: Frigorific Don Pedro, certificated (Livestock role 04.2.03.0002) by the Servicio Agrícola y Ganadero de Chile, and Frigorífico Cocarsa-Ecocarnes S.A., San Fernando, Provincia de Buenos Aires, Argentina, certified by the Servicio Nacional de Sanidad y Calidad Agroalimentaria de Argentina.

The data that support the findings of this study are available from the corresponding authors upon reasonable request.

## RESULTS

### Dynamin-dependent and dynamin-independent endocytosis induced by depolarization pulses in bovine adrenal chromaffin cells

In this study, we employed bovine chromaffin cells as our research model. These cells offer unique advantages for studying exocytosis and endocytosis with high temporal resolution, through electrophysiological and electrochemical techniques. Specifically, we focused on dynamin-2, a crucial isoform responsible for regulating exocytosis and endocytosis (González-Jamett et al., 2013a). In the results described below we sought to shed light on the effects of CNM-associated dynamin-2 mutations, A618T and S619L, on exocytosis and endocytosis in chromaffin cells.

Previously, we demonstrated in murine chromaffin cells that the application of short stimuli, particularly a single action potential-like stimulus (APls) or 5 ms square depolarization, activate a fast dynamin-dependent endocytosis, while longer stimuli (10 to 50 ms square depolarizations) induce a fast dynamin-independent endocytosis (Moya-Díaz 2016; Montenegro 2021; Bayonés 2022b). Here, we applied APls and 5 ms square depolarization to study the effect of CNM associated dynamin-2 mutations p.A618T and p.S619L on endocytosis in bovine chromaffin cells. As we mentioned, prior research was done in murine cells, so we first had to characterize the exocytotic and endocytotic responses elicited by the application of the same stimulation protocol, and their dependence on dynamin, in bovine chromaffin cells. Supplementary Figure 2 shows examples of Ca^2+^ currents (Fig. S2 A) and summarizes the Ca^2+^ currents amplitudes (I_Ca2+_) and integrals (òI_Ca2+_) (Fig. S2 B & C) obtained in response to the application of a APls and square depolarization pulses of 5 ms (named from now on as 5ms), respectively, in control conditions and in cells dialyzed with a monoclonal antibody against dynamin (Anti-Dyn). In addition, Figure 1 represents typical capacitance recordings obtained in response to the application of both types of depolarizations (Fig.1 A), and a summary of the different parameters associated to exocytosis and endocytosis obtained from the same experiments (Fig.1 B to E). In control conditions, similarly to murine cells (Bayonés et al., 2022b), bovine cells did not exhibit appreciable differences in ∫I_Ca2+_, exocytosis (ΔCm_exo_), or endocytosis (ΔCm_endo_). Additionally, also resembling murine cells, there were no effects of Anti-Dyn on I_Ca2+_, òI_Ca2+_, or ΔCm_exo_. More important, as shown in murine cells, ΔCm_endo_ and consequently endo/exo (which represents the efficiency of endocytosis computed as the ratio between ΔCm_endo_ and ΔCm_exo_) suffered a significant reduction when APls and 5ms were applied in the presence of Anti-Dyn (Figure 1 B, C & D). Lastly, as well as murine cells, the time constant of endocytosis (τ_endo_) was found to increased (indicating deceleration of endocytosis) in response to Anti-Dyn for APls and 5ms (Figure 1E). Overall, these findings provide valuable insights into the behavior of bovine cells in terms of fast endocytosis and its dependence with dynamin, confirming their similarity to what was described before in murine chromaffin cells.

**Figure 1.**
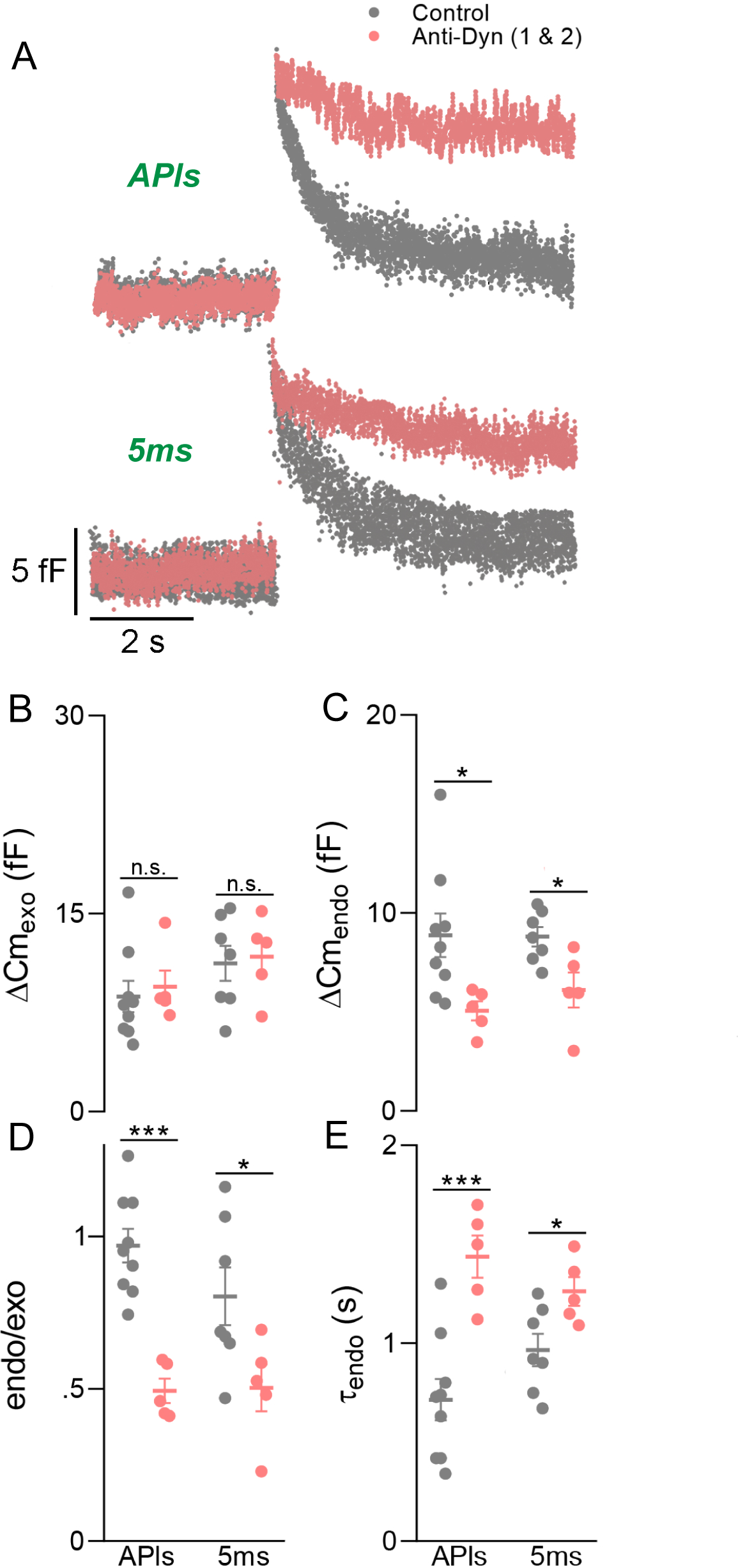
Anti-Dyn suppresses dynamin-dependent endocytosis evoked by action potential like-stimulus and short square depolarizations. (A) Representative examples of Cm recordings in response to the application of APls and 5ms stimuli obtained from two independent cells, where one was dialyzed with standard internal solution for 5 min (control, gray) and the other with a dynamin antibody (Anti-Dyn, red) also for 5 min. The plots in (B), (C), (D) and (E), represent the average values, standard errors, and individual measurements (one measurement/cell) of exocytosis (ΔCm_exo_), endocytosis (ΔCm_endo_), ratio between endocytosis and exocytosis (endo/exo), and time constant of endocytosis (τ_endo_), respectively, obtained by application of APls and 5ms square depolarizations in control (grey) and Anti-Dyn (red) conditions. Data were analyzed by Student’s ‘t’ test ((C): APls (t[12] = 2.479, p = 0.029), 5ms (t[10] = 2.874, p = 0.0166); (D): APls (t[12] = 5.883, p < 0.001), 5ms (t[10] = 2.303, p = 0.0441); (E): APls (t[12] = 4.466, p < 0.001), 5ms (t[10] = 2.574, p = 0.0277)). *p < 0.05, ***p < 0.001. These experiments were performed in 21 cells from 5 different cell cultures. The numbers between brackets represent the number of degrees of freedom handled in each comparison.

### Dynamin-2 p.A618T and p.S619L mutations inhibited dynamin-dependent endocytosis, but not dynamin-independent endocytosis, in adrenal bovine chromaffin cells

Subsequently, we proceeded to examine the impact of p.A618T and p.S619L mutations on endocytosis elicited by the aforementioned stimulation protocols. Figure 2A presents examples of capacitance measurements obtained from chromaffin cells expressing, separately, EGFP-wild-type (WT)-dynamin-2 or p.A618T or p.S619L, in response to the application of APls and 5ms. I_Ca2+_, òI_Ca2+_ (Fig.S3 B & C) and ΔCm_exo_ (Fig.2 B) were not affected by either of the mutants. Conversely, the expression of both mutants significantly decreased ΔCm_endo_ and the efficiency parameter endo/exo when the cells were stimulated by APls or 5ms (Fig 2C and D). Finally, the rate of endocytosis under the expression of any of both mutants increased their time constant τ (i.e., endocytosis was decelerated) for both type of stimuli (Fig 2E). In conclusion, the effects of p.A618T or p.S619L expression were comparable to the Anti-Dyn treatment, decreasing specifically the magnitude and rate of dynamin-dependent endocytosis induced by short depolarizations.

**Figure 2.**
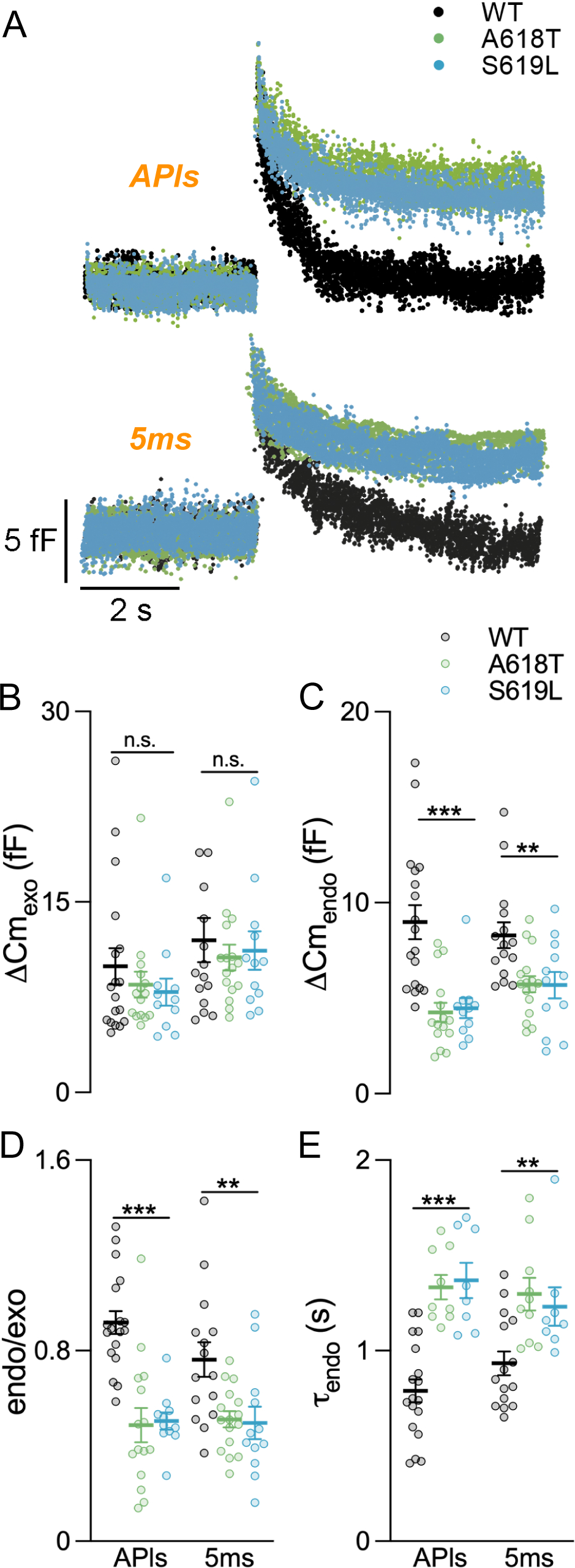
p.A618T and p.S619L affect dynamin-dependent endocytosis magnitude and dynamics. (A) Representative examples of Cm recordings in response to the application of APls obtained from three independent cells, where one expressed dynamin-2 WT (black), another A618T (green) and the other S619L (blue). The plots in (B), (C), (D) and (E), represent average values, standard errors, and individual measurements (one measurement/cell) of exocytosis (ΔCm_exo_), endocytosis (ΔCm_endo_), ratio between endocytosis and exocytosis (endo/exo), and time constant of endocytosis (τ_endo_), respectively, obtained by application of APls and 5ms square depolarizations, in WT (black), A618T (green) and S619L (blue), conditions. The data was analyzed by one-way ANOVA ((C): APls (F[2,41] = 14.39, p< 0.001), 5ms (F[2,40] = 6.432, p = 0.0038); (D): APls (F[2,41] = 20.12, p < 0.001), 5ms (F[2,40] = 6.524, p = 0.0035); (E): APls (F[2,32] = 23.36, p < 0.001), 5ms (F[2,30] = 6.778, p = 0.0037)). **p<0.01, ***p < 0.001. These experiments were performed in 46 cells from 7 different cell cultures. The numbers between brackets represent the degrees of freedom handled in each comparison.

### Dynamin-2 p.A618T and p.S619L mutations inhibited massive exo and endocytosis in adrenal bovine chromaffin cells

Next, we evaluated the effects of p.A618T and p.S619L on the exocytosis and endocytosis induced by a longer stimulus, such as a 500 ms square depolarization, which forces the mobilization and release of a bigger number of vesicles. Different authors found that such long depolarizations induce strong exocytosis followed by robust compensatory endocytosis or, alternatively, excess membrane retrieval (Engisch and Nowycky, 1998; Smith and Neher, 1997). In our hands, under the WT condition, this type of stimulation always induced a strong exocytosis followed by a slow compensatory endocytosis (Fig.3A). Notably, the expression of both mutants dramatically decreased the aforesaid parameters related with exocytosis (ΔCm_exo_), endocytosis (ΔCm_endo_) and endocytosis efficiency (endo/exo) (Fig.3 B, C and D). On the other hand, I_Ca2+_ and òI_Ca_2+ evoked by this stimulus were not affected by the expression of the mutants (Fig. S4 B, and C), which points towards that the observed reductions in exocytosis and endocytosis are not associated with alterations in calcium entry. The almost complete abolishment of endocytosis in cells transfected with p.A618T and p.S619L suggest that this process is associated to a dynamin-dependent mechanism, as was suggested by previous studies (Engisch and Nowycky, 1998). In addition, the huge inhibition of ΔCm_exo_ observed in these conditions, together with the lack of effect of p.A618T and p.S619L on this parameter for short depolatizations, suggest that both mutants affect a process that becomes important when exocytosis exceeds a certain number of vesicles.

**Figure 3.**
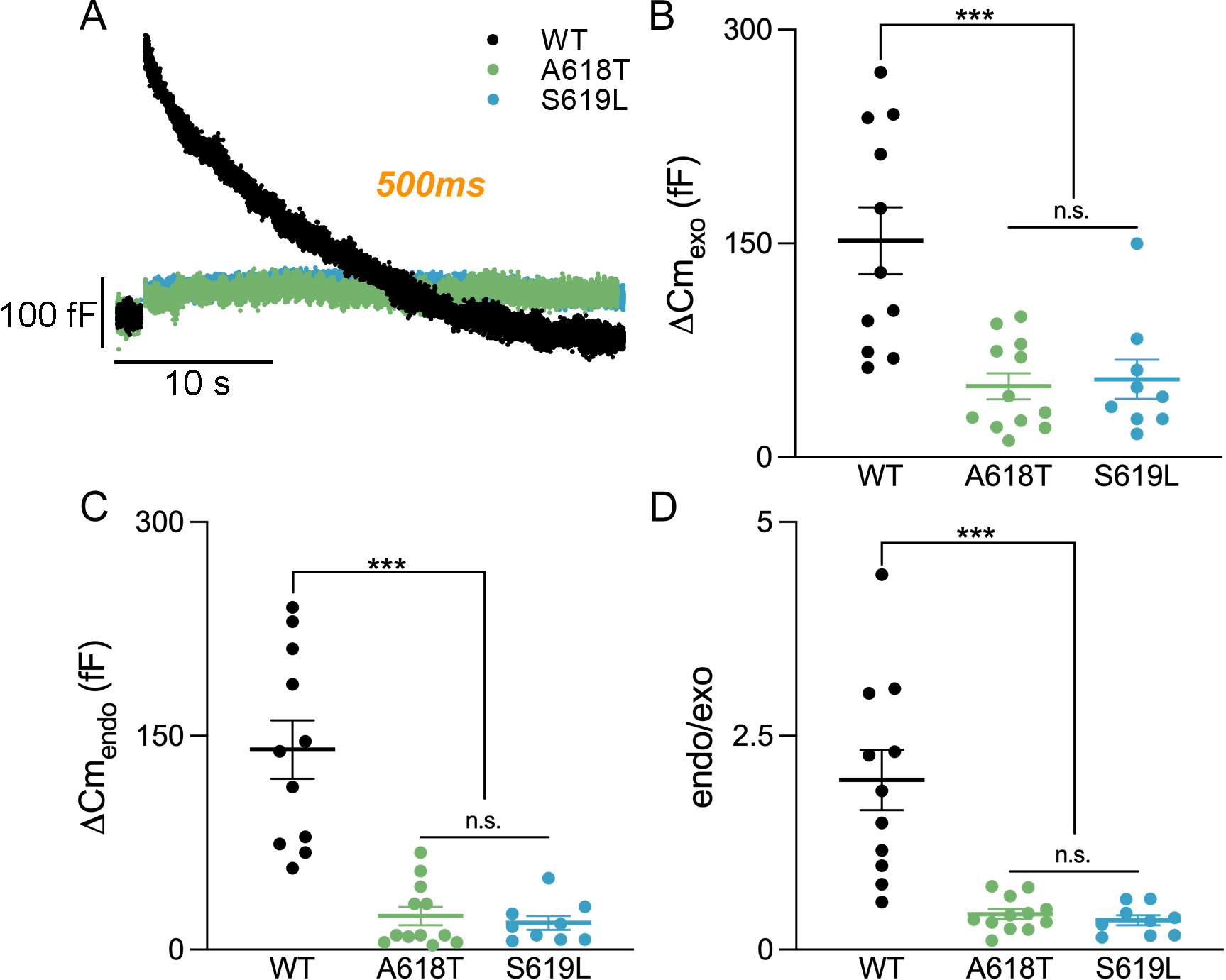
Dynamin-2 mutations A618T and S619L disrupts exocytosis and endocytosis evoked by prolonged stimulation. (A) Representative examples of Cm recordings in response to the application of a 500 ms square depolarization obtained from three independent cells, where one expressed dynamin-2 WT (black), another A618T (green) and the other S619L (blue). The plots in (B), (C) and (D) represent average values, standard errors, and individual measurements (one measurement/cell) of exocytosis (ΔCm_exo_), endocytosis (ΔCm_endo_) and ratio between endocytosis and exocytosis (endo/exo), respectively, obtained by application of 500 ms square depolarization, in WT (black), A618T (green) and S619L (blue), conditions. The data was analyzed by one-way ANOVA ((B): F[2,29] = 12.39, p < 0.001); (C): F[2,29] = 27.83, p< 0.001); (D): F[2,29] = 18.78, p < 0.001)). ***p < 0.001. These experiments were performed in 31 cells from 5 different cell cultures. The numbers between brackets represent the degrees of freedom handled in each comparison.

### Dynamin-2 p.A618T and p.S619L Mutations Reduce the Secretion induced by a nicotinic agonist in adrenal bovine chromaffin cells

Bovine chromaffin cells constitute a classical experimental model to study neurosecretion, a cellular process that is also influenced by dynamin-2 (González-Jamett et al., 2013a). Therefore, to analyze the effects of the p.A618T and p.S619L on exocytosis using a different technique and a more physiological approach, we evaluated the release of catecholamine induced with the activation of nAChRs using carbon fiber amperometry. The exocytosis was evoked by a 10 s pulse of the nAChRs agonist DMPP applied to chromaffin cells expressing alternatively EGFP-WT-dynamin-2, or A618T, or S619L. Examples of typical recordings are represented in Figure 4A. As shown in Figure 4B, the number of DMPP-induced exocytotic events was significantly reduced by both mutations, as compared with those expressing EGFP-WT-dynamin-2.

**Figure 4.**
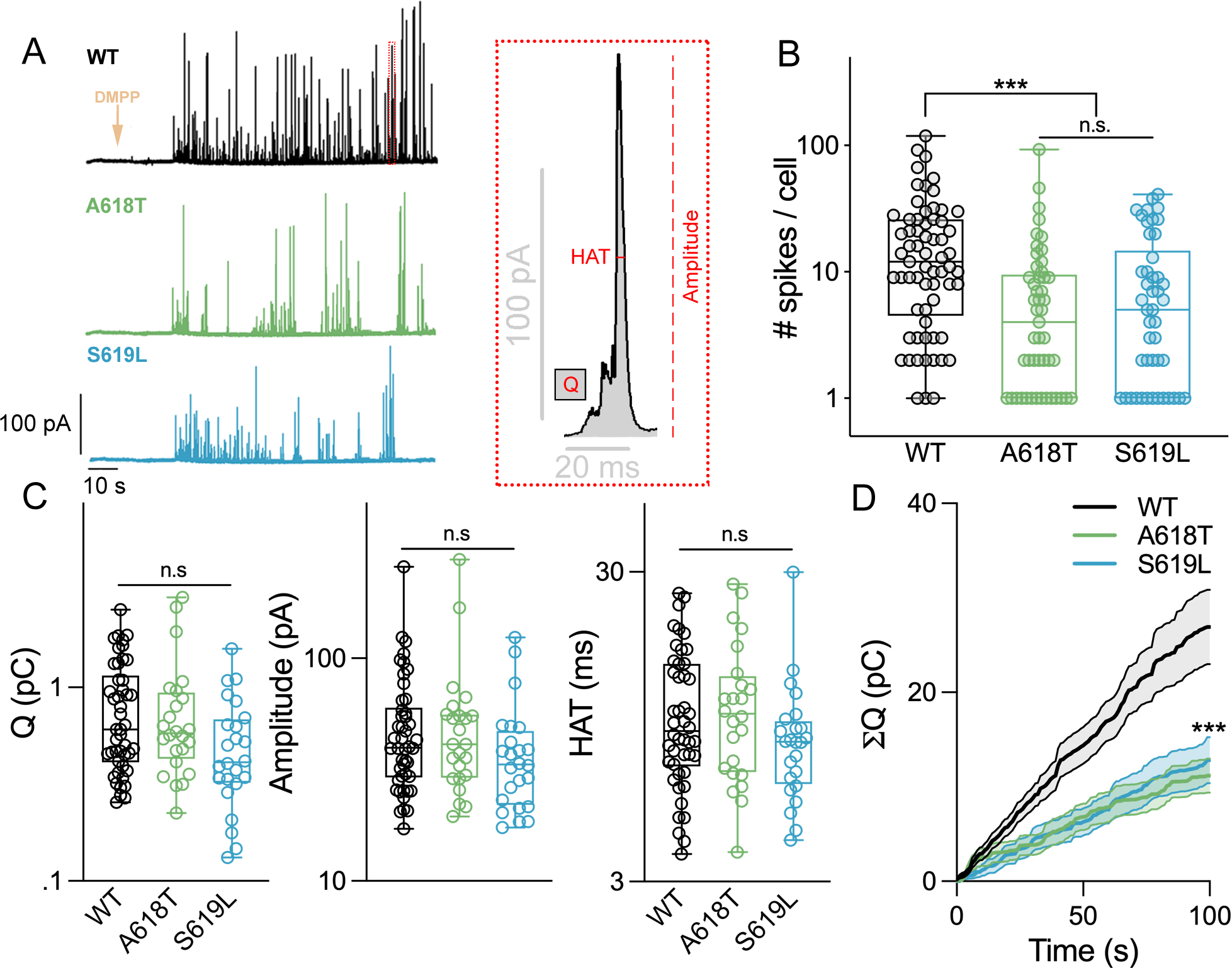
Dynamin-2 mutations disrupt secretion provoked by nicotinic agonist long-lasting stimulation. (A) Left: Representative amperometric recordings obtained from three individual cells stimulated with the nicotinic agonist DMPP for 10 s (orange arrow), one expressing dynamin-2 WT (black), another A618T (green) and the other S619L (blue). Right: a representative amperometric spike, displaying a pre-spike foot signal (PFS), extracted from the WT trace on the left (see reference dotted red box). The parameters Q (area under the spike), amplitude (length of the spike from the basal line) and HAT (elapsed time at spikés half amplitude) are marked in red. (B) The number of spikes per cell (each colored empty circle) is displayed for the three conditions, WT (black), A618T (green) and S619L (blue), in a log10 scale. Cells expressing the mutations p.A618T and p.S619L significantly decreased its response when compared against the WT condition (C) The parameters Q, amplitude, and HAT are displayed in the three conditions, WT (black), A618T (green) and S619L (blue), in a log10 scale. Each empty circle represents the median obtained from the totality of spikes recorded in each cell. Neither of these parameters are significantly changed between conditions. For (B) and (C), the centered, inferior, and superior lines in the box plot represents the median, and first and third quartiles, respectively. Top and bottom whiskers lines show the maximum and minimum values of each distribution, respectively. (D) Cumulative integration of the Q parameter (ΣQ) for each experimental condition (WT, A618T and S619L). Each individual experiment (one measurement per cell) was binned in intervals of 1 s, all experiments were averaged, and represented as means ± SEM. ΣQ decreased markedly in cells transfected with A618T or S619L. In B and C data was analyzed by using a Kruskal-Wallis test followed by Dunn’s multiple comparison test against WT condition, while in D a Kolmogorov-Smirnov test was applied. ***p < 0.001. These experiments were performed in 96 cells from 7 different cell cultures.

Amperometry also allows the analysis of the dynamics of catecholamines release from individual exocytosis events (Mosharov and Sulzer, 2005). To determine whether A618T or S619L affect the characteristics of the individual exocytotic events, we further analyzed different spike parameters (cartoon on Fig. 4 A), particularly the spike charge (Q), that is proportional to the amount of catecholamines released during each individual event, the spike amplitude, which accounts for the height of the spike, and time elapsed at spikés half amplitude (HAT). The latter is related to the duration of the release event through an expanded fusion pore (Fang et al., 2008). These parameters where not modified by the expression of A618T or S619L mutations when compared against WT (Fig. 4 C). As the mutations did not modified Q but reduced the number of DMPP-induced exocytotic events, the cumulative Q along experimental time (ΣQ), which is a measurement of the total catecholamine release, was markedly reduced by both mutants (Fig. 4D).

In addition, we also analyzed the amplitude and the duration of the pre-spike foot signal (PSF; Fig. S5 B), which respectively reflect the conductance and stability of the fusion pore (Lindau and Alvarez de Toledo, 2003). The fusion pore is an unstable channel formed during the fusion process (Marengo and Cárdenas, 2017). As shown in the supplementary Figure 5, C and D, neither A618T nor S619L mutation significantly changed these fusion pore parameters.

### Dynamin-2 p.A618T and p.S619L Mutations Reduce cortical NPY labeled vesicles in adrenal bovine chromaffin cells

A possible explanation for the effect of both mutations on exocytosis might be the reduction of vesicles available for fusion. It is well known that dynamin activity modulates actin dynamics (Zhang et al., 2020), and actin is involved in the transport of vesicles to plasma membrane (Gimenez-Molina et al., 2018). In consequence, if the expression of p.A618T and p.S619L affect actin dynamics, it is possible that these mutations may disrupt the trafficking of vesicles to the plasma membrane. To assess this hypothesis, we co-expressed NPY- mCherry together with WT dynamin, or p.A618T or p.S619L, fixed the cells, and evaluated the area covered by NPY associated fluorescence by confocal microscopy (see Materials and Methods for more details). It is important to mentioned that NPY is normally localized in the core of secretory vesicles in physiological conditions (Hwang et al., 2007). We performed these experiments in control rest conditions or after stimulating the cells with 50 μM de DMPP for 10 s before fixation.

Figure 5 (top) show examples of confocal images of cells expressing WT, p.A618T or p.S619L in unstimulated (Fig. 5A) and stimulated conditions (Fig. 5B). It is possible to observe the distribution of NPY (red) throughout the cytosol, and particularly in localized spots in the cell border, probably representing vesicles or vesicle clusters. Dynamin-2 constructs (green) were also distributed throughout the cytosol, and for both mutants it is possible to note the accumulation of fluorescence in big spots, possibly representing mutated dynamin polymers as it was described previously (Wang et al, 2010). The bottom of Figure 5 displays the quantification of the area covered by NPY associated fluorescence in the whole cell (% NPY Total) and in a ring of 1 µm from the cell border (% NPY Periphery), and the ratio between these two areas (% Periphery/Total). The expression of both mutants did not alter any of these parameters in unstimulated cells. However, both mutants decreased very significantly the NPY in the periphery of stimulated cell. These results indicate that the expression of p.A618T or p.S619L impairs the recruitment of secretory vesicles to the cell surface after exocytosis-induced stimulus.

**Figure 5.**
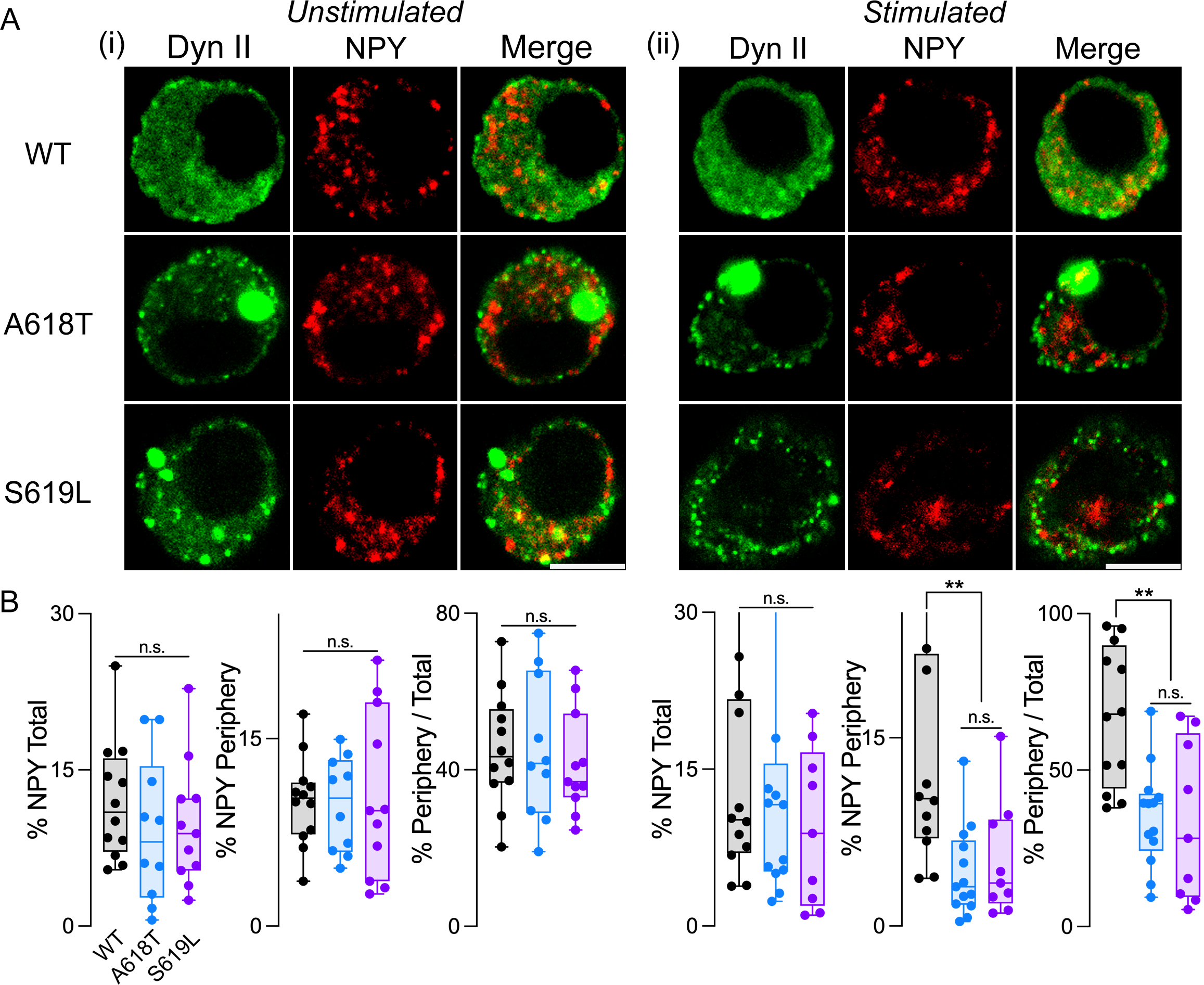
Dynamin-2 mutations affect vesicle translocation to the plasma membrane in stimulated chromaffin cells. (A) Representative images, taken at the equatorial plane, of three unstimulated (I) or DMPP-stimulated (II) chromaffin cells co-expressing separately the Dyn-2 constructs EGPF-tagged WT (top row), p.A618T (middle row) or p.S619L (bottom row) and the vesicle fluorescent marker NPY-mCherry. Stimulation with 50 µM DMPP lasted 10 s. EGFP fluorescence is represented in the left column (Dyn II), mCherry fluorescence is represented in the middle column (NPY), while the right column depicts the merge of both fluorescence channels. Scale bar = 5 μm. (B) Percentage of the area occupied by NPY-Cherry with respect to the celĺs full area (% NPY Total), or percentage of the area occupied by NPY- Cherry in a peripheral area of 1 μm wide aligned with the plasma membrane (% NPY Periphery), or the ratio % NPY Periphery/% NPY Total, in unstimulated (I) or DMPP stimulated (II) chromaffin cells expressing Dyn-2 WT (black), p.A618T (blue) and p.S619L (violet). Each cell is represented by filled colored dots, while the centered, inferior and superior lines in the box plot represents the median, first and third quartile, respectively. Top and bottom whiskers lines show the maximum and minimum values of each distribution, respectively. In unstimulated cells, no differences were found among the three conditions for all calculated parameters. On the other hand, for stimulated cells, %NPY Periphery and the ratio % Periphery/Total drastically decreased for those cells expressing the mutations of Dyn-2. Data in panels A and B was analyzed by using a Kruskal-Wallis test followed by Dunn’s multiple comparison test against WT condition. **p < 0.01. These experiments were performed in 33 cells from 3 different cell cultures.

### Dynamin-2 p.A618T and p.S619L mutations disturb F-actin dynamics

It is well known that endocytosis and exocytosis are influenced by actin dynamics (Grassart et al., 2014; Gabel et al., 2015; Cárdenas et al., 2022; Montenegro et al, 2021), and that A618T and S619L mutations disrupt the formation of actin filaments (F-actin) in skeletal myoblasts (Bayonés et al., 2022a). In consequence, we decided to analyze the effect of these mutations on actin dynamics in bovine chromaffin cells. With this goal in mind, we transfected chromaffin cells with mCherry WT, or A618T or S619L dynamin-2. F-actin *de novo* polymerization was evaluated 48 h later by incubating permeabilized chromaffin cells in presence of Alexa Fluor 488-tagged actin monomers (see Materials and Methods for the details of the incubation solution). In this assay, the fluorescence signal corresponds to the recently formed actin filaments (González-Jamett et al., 2013a; González-Jamett et al., 2017); therefore, the percentage of F-actin formation was computed as a ratio between the area occupied by fluorescent F-actin and celĺs total area (see Methods for further details). As shown in the examples represented in Figure 6A and in the quantification of Figure 6B, both dynamin-2 mutations significantly reduced the formation of new actin filaments.

**Figure 6.**
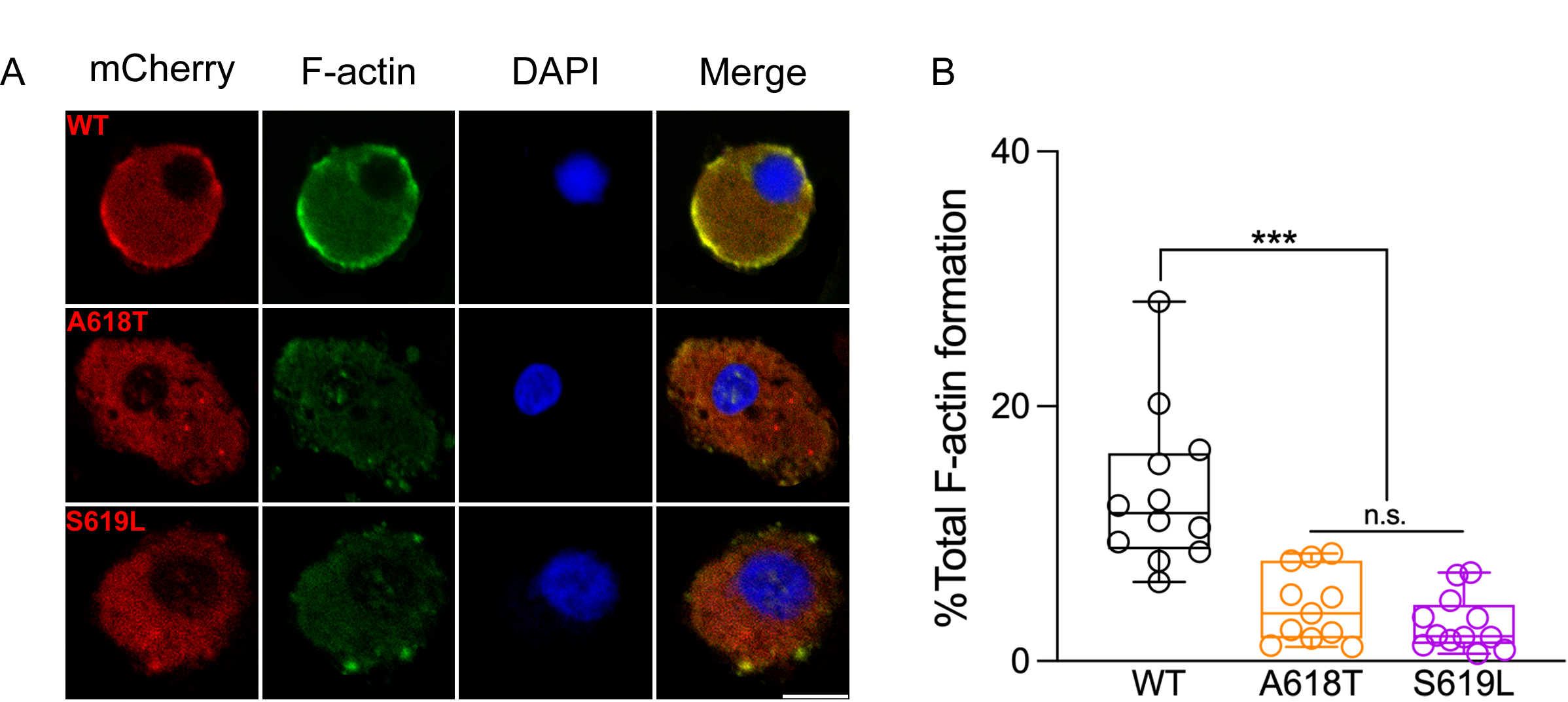
Dynamin-2 mutations impaired *de novo* F-actin formation in chromaffin cells. (A) Representative images of three independent chromaffin cells expressing Dyn-2 WT (top row), or p.A618T (middle row) or p.S619L (bottom row). The columns show (from left to right) the expression of Dyn-2 fused to the mCherry fluorescent marker (mCherry column); the F-actin *de novo* formation, assessed by incubating the transfected cells with a G-actin Alexa Fluor 488 conjugate (F-actin column); the nucleus, stained with DAPI fluorescent probe (DAPI column); and the merging of the three previous channels. Scale bar = 5 μm. (B) The F-actin *de novo* formation per cell (empty colored circles), computed as the ratio (%) between the area occupied by Alexa Fluor 488 F-actin fluorescence and celĺs full area (see Methods), are shown for WT (black), A618T (orange) and S619L (pink). The centered, inferior and superior lines in the box plot represents the median, first and third quartile, respectively. Top and bottom whiskers lines show the maximum and minimum values of each distribution, respectively. The mutations p.A618T and p.S619L notably decreased the F-actin *de novo* formation when compared against the WT condition. Data in B was analyzed by using a Kruskal-Wallis test followed by Dunn’s multiple comparison test against WT condition. ***p < 0.001. These experiments were performed in 35 cells from 4 different cell cultures.

## DISCUSSION

Dynamin-2 was classically associated to endocytosis, however it has also been implicated in other cellular processes, like vesicle trafficking, exocytosis, and regulation of actin filament dynamics (González-Jamett et al., 2013b). Recently, by using total internal reflection fluorescence microscopy in immortalized human myoblasts, we reported that p.A618T and p.S619L dynamin-2 mutations impairs Ca^2+^-induced exocytosis of GLUT4 containing vesicles (Bayonés et al, 2022a). These two mutations have been linked to severe neonatal forms of CNM (Böhm et al., 2012), a form of skeletal muscle hereditary disease. However, since dynamin-2 is ubiquitously present in a wide variety of tissues (Cao et al, 1998), it is possible that these mutations, which cause a severe neonatal phenotype, may also affect other cell types, including neuroendocrine cells and neurons. In this work we analyze if the CNM associated mutations p.A618T and p.S619L alter exocytosis and endocytosis in chromaffin cells, a classical model for the study of neurosecretion.

Our study shows that both mutations inhibit specifically: (1) a form of dynamin dependent endocytosis that is activated after the application of brief single depolarizations (Bayonés et al, 2022b); (2) the exocytosis and endocytosis triggered by prolonged 500 ms depolarizations; (3) the catecholamine release triggered by the nicotinic pathway activation; and (4) NPY labeled vesicle recruitment to the cell cortex; and finally (5) the *de novo* formation of F-actin filaments.

We wondered if there is a common factor that might explain so many processes affected by these dynamin-2-mutants. It was described in numerous publications that alterations in actin dynamics affect both exocytosis and endocytosis in different systems (Miklavc and Frick, 2020; Wu and Chan, 2022). In this regard, it was shown by several authors, starting three decades ago, that interfering in actin polymerization/depolymerization dynamics affect secretory vesicles transport to plasma membrane, and consequently exocytosis (Trifaró et al., 2002; Meunier and Gutiérrez, 2016; Gutiérrez and Villanueva, 2018). Some investigators have shown that actin filaments are also involved in the mechanism of fusion itself (Doreian et al., 2008; Berberian et al., 2009; Wen et al., 2016). Finally, it was demonstrated that actin filaments also participate in several endocytosis mechanisms, as bulk endocytosis, classical clathrin-mediated endocytosis, and ultrafast endocytosis (Boulant et al., 2011; Watanabe et al., 2013; Gormal et al., 2015). Particularly, we showed recently that the dynamin-dependent endocytosis induced after short depolarizations, like the one shown in the first section of this paper, is also dependent on polymerized F-actin (Montenegro et al. 2021). In the present work, we observed that p.A618T and p.S619L mutations disrupt the formation of actin filaments in chromaffin cells, and very recently we described a similar effect of both mutations in immortalized human myoblasts (Bayonés et al, 2022a). It is important to note that, from the beginning, dynamin was associated with actin dynamics (Gu et al., 2010; Zhang et al., 2020). It is therefore possible that the disruption of actin filaments formation might be responsible of the effects of these dynamin-2 mutation on endocytosis and neurosecretion.

Other dynamin-2 mutations that cause CNM, such as the mutants R369W and R465W, or mutations linked to Charcot-Marie-Tooth neuropathy like the K562E, also lead to aberrant actin dynamics (Yamada et al., 2016b; González-Jamett et al., 2017; Hamasaki et al. 2022; Arriagada-Diaz et al., 2023). Interestingly, the R465W mutation further perturbs the synaptic plasticity and cognitive function in a mouse model of CNM (Arriagada-Diaz et al., 2023), supporting the pleiotropic role of these mutations. Why these mutations that seem to display increased GTPase activity (Kenniston and Lemmon, 2010) disrupt actin dynamics? A possible explanation is that the enhanced self-assemble of the CNM mutations (Kenniston and Lemmon, 2010; Wang et al., 2010) that lead to the accumulation of large oligomers in the cytosol (James et al., 2014; González-Jamett et al., 2017) renders dynamin less able to interact with F-actin upon a stimulus. The augmented self-assemble of the CNM mutations can also result in the formation of abnormal clathrin-containing structures at the plasma membrane (James et al., 2014) with altered internalization (Ali et al., 2019).

The intricate functions of dynamins, particularly dynamin-2, and the detrimental consequences of disease-associated mutations underscore the critical role of these proteins in cellular homeostasis and disease pathogenesis. Thus, by elucidating the impact of specific mutations on endocytosis, neurosecretion, and cytoskeletal dynamics, this work shed light on the underlying mechanisms and provide valuable insights into the pathophysiology of centronuclear myopathy.

## Acknowledgments

This work was supported by the grants PICT 2764-2016, PICT 02849-2018 and PICT I-A- 00176 - 2021 (to FDM) from the Agencia Nacional de Promoción Científica y Tecnológica (Argentina), and FONDECYT 1220825 (to AMC) from ANID (Chile). LB has now a postdoctoral fellowship (# 838783) from CONAHCYT México. We thank CONAHCYT, and Dr. Román Rossi Pool for their positive cooperation during the analysis of data and confection of this manuscript. We also thank to MSc Sergio Parra for helping us with the analysis of amperometry experiments.

## Author Contributions

LB performed experiments, analyzed data, designed and prepared the figures, participated in the writing and revision of the manuscript; MJGF performed experiments and analyzed data, SAB performed experiments; LIG designed and performed analysis of NPY images; CFC performed experiments; XBM performed experiments; AGJ designed experiments and edited the manuscript; AMC conceived the study, designed experiments, and drafted the manuscript; FDM conceived the study, designed experiments, and drafted the manuscript. All authors contributed to interpretation of data and revised the final version of the manuscript. All authors have read and agreed to the published version of the manuscript.

