## Supplementary Material for "Dynamin-2 Mutations Linked to Neonatal-onset Centronuclear Myopathy impair exocytosis and endocytosis in adrenal chromaffin cells"

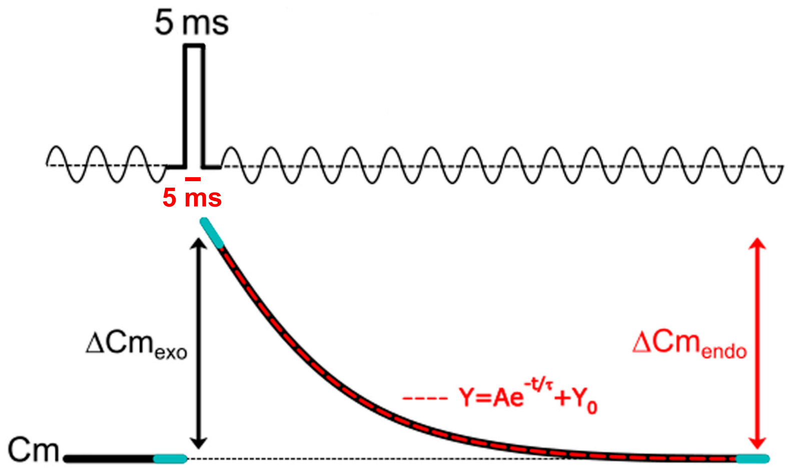


**Supplementary Figure 1.** This cartoon outlines a 5 ms square voltage pulse (top) and the resulting change in membrane capacitance (bottom). The light blue lines on the capacitance recording scheme represent the segments used to estimate the Cm values at the initial baseline, at the peak of Cm increase after the pulse and at the end of Cm decay, from where $\Delta Cm_{exo}$ and $\Delta Cm_{endo}$ were estimated. The dotted red line represents the portion of the capacitance recording scheme used for the single exponential fitting of endocytosis. This analysis can be generalized to other stimulation pulses applied in this study.

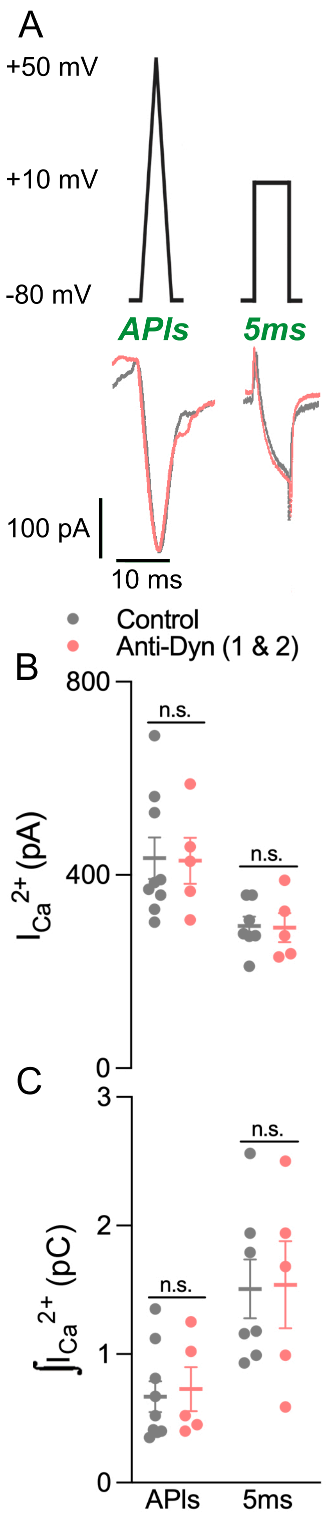


**Supplementary Figure 2.** (A) Illustration of an APIs and a 5 ms square pulse (top) and original recordings of calcium currents (bottom) obtained from two independent cells, one in control conditions (Control, grey) and the other with the application of a monoclonal anti-dynamin antibody (7 nM) in the internal solution (Anti-Dyn, red). (B)-(C) The plots represent average values, standard errors and individual measurements (one measurement per cell) of $I_{Ca2+}$ and $\int I_{Ca2+}$ obtained by application of APIs and 5 ms square pulse in control condition (grey circles, n=9 and n=7, respectively) and with Anti-Dyn (red circles, n=5 and n=5, respectively). The data were analyzed by Student's 't' test. No difference was found between conditions.

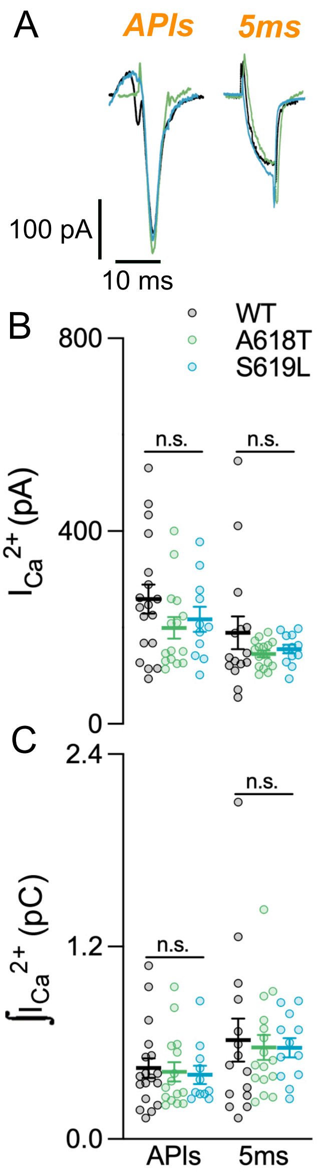


**Supplementary Figure 3.** (A) Representative recordings of $Ca^{2+}$ currents evoked by the application of an APls or a 5 ms square pulse obtained from three independent cells, one expressing Dyn-2 WT (WT, black), other the mutation p.A618T (A618T, blue) and another the mutant p.S619L (S619L, green). (B)-(C) The plots represent average values, standard errors and individual measurements (one measurement per cell) of $I_{Ca2+}$ and $\int I_{Ca2+}$ obtained by application of APls and 5 ms square pulse in cells expressing Dyn-2 WT (black open circles, n=18 and n=15, respectively), A618T (green open circles, n=15 and n=16, respectively), and S619L (blue open circles, n=11 and n=12, respectively). The data was analyzed by one-way ANOVA followed by a Bonferroni´s comparison test against WT condition. No difference was found between conditions.

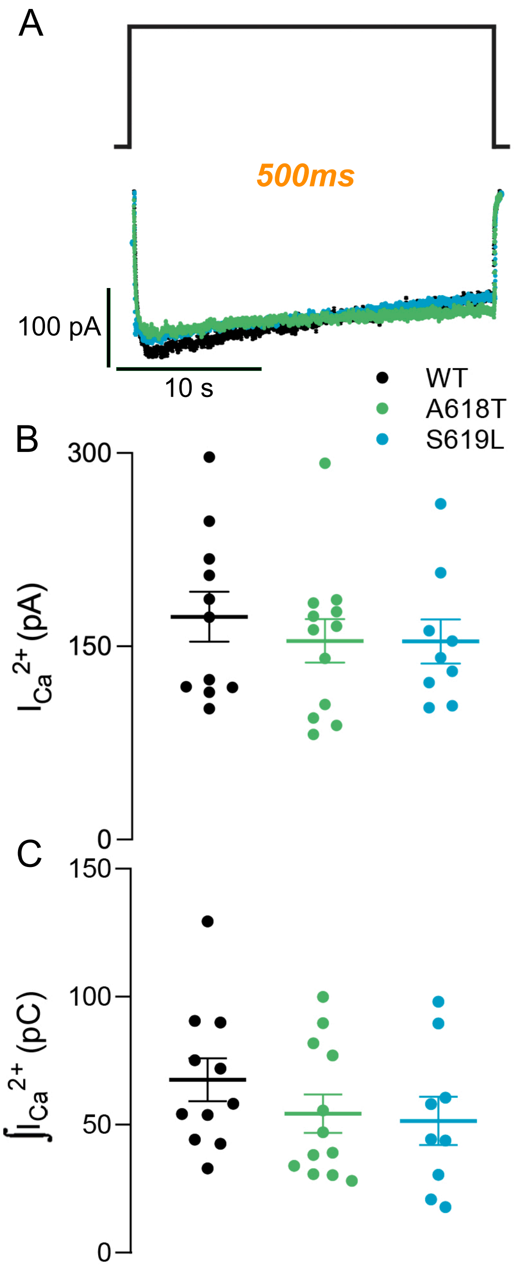


**Supplementary Figure 4.** (A) Illustration of a 500 ms square pulse (top) and original representative recordings of $Ca^{2+}$ currents (bottom) obtained from three independent cells, expressing Dyn-2 WT (WT, black), or the mutation p.A618T (A618T, blue) or p.S619L (S619L, green). (B)-(C) The plots represent average values, standard errors and individual measurements (one measurement per cell) of $I_{Ca2+}$ and $\int I_{Ca2+}$ obtained by application of 500 ms square depolarizations in cells expressing Dyn-2 WT (black circles, n=11), A618T (green circles, n=12), and S619L (blue circles, n=9). The data was analyzed by one-way ANOVA followed by a Bonferroni´s comparison test against WT condition. No difference was found between conditions.

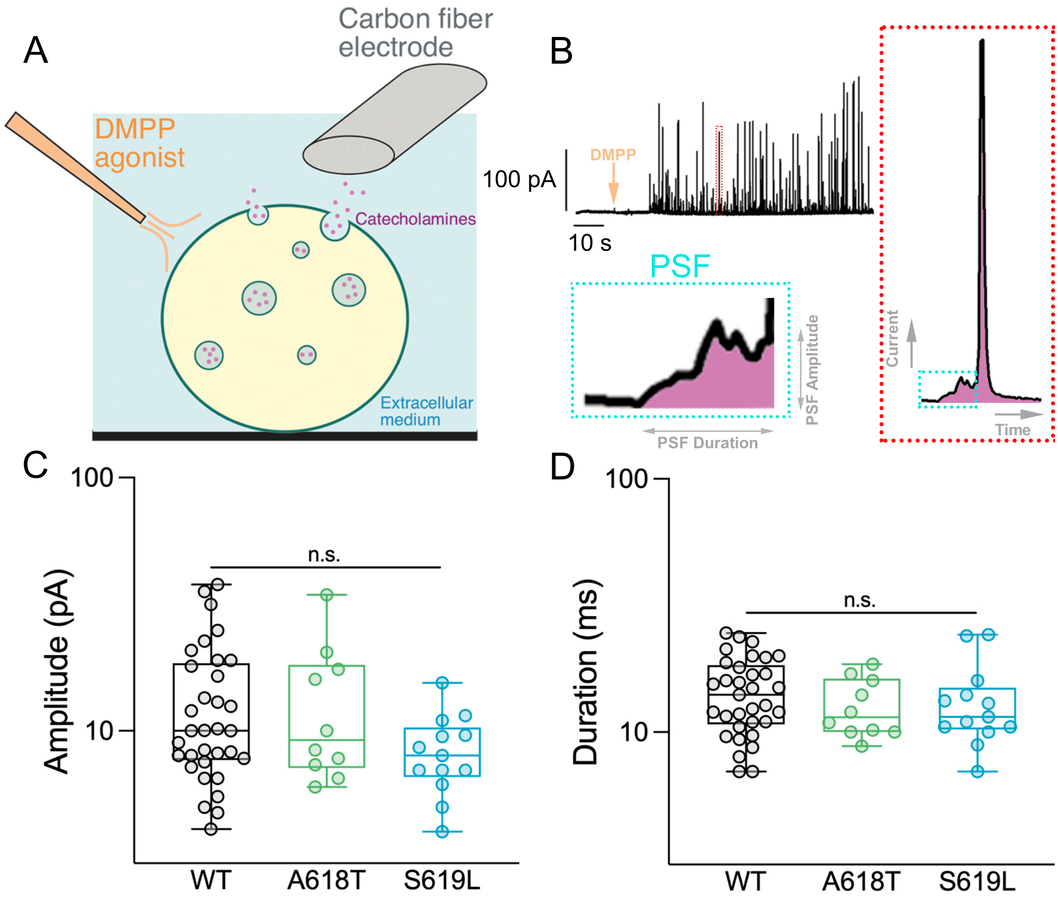


**Supplementary Figure 5.** (A) The cartoon illustrates the experimental paradigm used for the obtention of amperometric data. A carbon fiber electrode (on the right) is placed softly on the surface of a chromaffin cell, while a stimulation micropipette loaded with DMPP (on the left) is approximately 3-5 cell diameters from the cell. For clarity, these distances are not represented in the cartoon. (B) Top Left, a typical amperometric recording in a WT cell. After a baseline of 10 s was obtained, a DMPP puff of 10 s (the arrow indicates the beginning of the puff) was applied, then the recording continued for another additional 80 s. Right, magnified amperometric spike (red dotted box) showing a typical pre-spike foot signal (PSF) (cyan dotted box). Bottom Left, magnified PSF of the spike represented at the right, from where the amplitude and duration are calculated. (C)-(D) The amplitude and duration of the PSF per cell (each colored circle) is displayed for the three conditions, WT (black, n=33), A618T (green, n=10) and S619L (blue, n=13), in a log10 scale. The centered, inferior, and superior lines in the box plot represents the median, and first and third quartile, respectively. Top and bottom whiskers lines show the maximum and minimum values of each distribution, respectively. Neither of these parameters significantly changed between conditions. The data were analyzed by using a Kruskal-Wallis test followed by Dunn’s multiple comparison test against WT condition.
